# ArcheD, a residual neural network for prediction of cerebrospinal fluid amyloid-beta from amyloid PET images

**DOI:** 10.1101/2023.06.20.545686

**Authors:** Arina A. Tagmazian, Claudia Schwarz, Catharina Lange, Esa Pitkänen, Eero Vuoksimaa, the Alzheimer’s Disease Neuroimaging Initiative

## Abstract

Detection and measurement of amyloid-beta (Aβ) aggregation in the brain is a key factor for early identification and diagnosis of Alzheimer’s disease (AD). We aimed to develop a deep learning model to predict Aβ cerebrospinal fluid (CSF) concentration directly from amyloid PET images, independent of tracers, brain reference regions or preselected regions of interest. We used 1870 Aβ PET images and CSF measurements to train and validate a convolutional neural network (“ArcheD”). We evaluated the ArcheD performance in relation to episodic memory and the standardized uptake value ratio (SUVR) of cortical Aβ. We also compared the brain region’s relevance for the model’s CSF prediction within clinical-based and biological-based classifications. ArcheD-predicted Aβ CSF values correlated strongly with measured Aβ CSF values (*r*=0.81; *p*<0.001) and showed correlations with SUVR and episodic memory measures in all participants except in those with AD. For both clinical and biological classifications, cerebral white matter significantly contributed to CSF prediction (*q*<0.01), specifically in non-symptomatic and early stages of AD. However, in late-stage disease, brain stem, subcortical areas, cortical lobes, limbic lobe, and basal forebrain made more significant contributions (q<0.01). Considering cortical gray matter separately, the parietal lobe was the strongest predictor of CSF amyloid levels in those with prodromal or early AD, while the temporal lobe played a more crucial role for those with AD. In summary, ArcheD reliably predicted Aβ CSF concentration from Aβ PET scans, offering potential clinical utility for Aβ level determination and early AD detection.

## 1. Introduction

Early detection and diagnosis of Alzheimer’s disease (AD) can help in the prevention of dementia and in identifying at-risk individuals for clinical drug or lifestyle intervention trials. Even though AD is characterized by both clinical symptoms and neuropathological changes, diagnostic guidelines have been based on clinical symptoms, in particular on episodic memory impairment (Jack et al. 2011; McKhann et al. 1984). However, clinical classification based on symptoms is neither sensitive nor specific for AD, as cognitive impairment can be caused by other neurodegenerative diseases or other reasons (Erickson et al. 2022; Jack et al. 2018; Sperling, Karlawish, and Johnson 2013). Approximately 10-30% of those with clinically defined AD do not have AD-specific neuropathological changes of amyloid-beta (Aβ) plaques and neurofibrillary tangles of hyperphosphorylated tau on autopsy (Nelson et al. 2011). Moreover, about 30% of cognitively healthy individuals have substantial AD-related neuropathological changes (Aizenstein et al. 2008; Knopman et al. 2003).

Acknowledging the inconsistency between cognitive status and AD-related brain pathology, the biological classification of AD is based solely on biomarkers and defines AD independently of the cognitive status (Jack et al. 2011). The AT(N) framework – and its extension ATX(N) – are based on biological hallmarks of AD: Aβ (A), tau (T) related pathology, and also neurodegeneration (N) (Jack et al. 2018) and potential novel biomarkers (X) (Hampel, Cummings, et al. 2021). Aggregation of Aβ can start even decades before the onset of first clinical symptoms, whereas tau protein deposition begins much later and closer to first clinical symptoms (Jack et al. 2010). Based on Aβ and tau measurements in cerebrospinal fluid (CSF) and positron emission tomography (PET) and determination of neurodegeneration via PET or magnetic resonance imaging (MRI) scans, the AT(N) framework yields eight biomarker profiles which separate the AD continuum from non-AD pathologic changes and normal AD biomarkers (Jack et al. 2018). In the current framework, it is the Aβ positivity that is decisive and defines individuals to be in the AD continuum (Jack et al. 2018). Further, cognitive status is evaluated independently of the biomarker profile and is used for disease staging. Although biomarkers are currently supplementary in clinical practice of AD definition (Jack et al. 2011) and/or limited to memory clinics, biological classification is intended to improve the definition, classification, and diagnosis of AD as a unique neurodegenerative disease (Hampel et al. 2022).

Large neuroimaging datasets with PET measurements of AD biomarkers provide an excellent opportunity to further evaluate the AT(N) framework (Weber et al. 2021). Deep learning (DL) models have shown high accuracy in classifying AD and its progression from MRI and PET images (Jo, Nho, and Saykin 2019). While the majority of these models have used MRI or PET scans to predict clinical disease stages, early disease detection, or disease progression (Choi et al. 2020; Ding et al. 2019; Jo et al. 2020; Lin, Lin, and Lane 2021), only a few studies have aimed to predict AD fluid biomarkers based on PET (Kim et al. 2019; F. Reith et al. 2020; F. H. Reith, Mormino, and Zaharchuk 2021). So far, models have not been trained to predict biological classes defined by amyloid CSF or the biomarker values by itself directly from the amyloid PET, independent of the Aβ tracer type, preselected regions of interest, brain reference region and standardized uptake value ratio (SUVR) calculation (Spallazzi et al. 2019; Palmqvist et al. 2015). Such a prediction is needed to study the link between the brain and CSF amyloid and will also benefit work on blood-based biomarkers of AD.

In this work, we present a novel approach of predicting the Aβ CSF concentration from amyloid PET images with a deep neural network model (“ArcheD”) based on convolutional residual networks. Our approach allows probing the model and input amyloid PET images to reveal the brain regions that contribute most to the predicted CSF amyloid concentrations. ArcheD does not use any prespecified cortical (or other) brain regions, but instead adopts a hypothesis-free approach leveraging all information available in PET images. To examine our approach, we compared our method’s performance with SUVR, a measure that determines cortical amyloid aggregation in relation to cerebellum, a commonly used reference region (Heeman et al. 2020; Jack et al. 2013). We also studied our model in relation to episodic memory including immediate and delayed recall measures. We further investigated the model and brain regions separately in sub-groups based on both clinical and biological classification of AD. To scrutinize the trained neural network model, we extracted brain regions which the model considers informative for CSF prediction and compared them between clinical and biological classes.

## 2. Methods and materials

### 2.1. Data and participants

We studied 1252 individuals’ PET data on brain amyloid and CSF measurements of amyloid-β 1-42 peptide and phosphorylated tau 181P provided by the Alzheimer’s Disease Neuroimaging Initiative (ADNI).

Data used in the preparation of this article were obtained from the Alzheimer’s Disease Neuroimaging Initiative (ADNI) database (adni.loni.usc.edu). The ADNI was launched in 2003 as a public-private partnership, led by Principal Investigator Michael W. Weiner, MD. The primary goal of ADNI has been to test whether serial magnetic resonance imaging (MRI), positron emission tomography (PET), other biological markers, and clinical and neuropsychological assessment can be combined to measure the progression of mild cognitive impairment (MCI) and early Alzheimer’s disease (AD). (Petersen et al. 2010)

We used PET and MRI scans, CSF measurements, ADNI clinical classifications, episodic memory measurements and demographics data (adni.loni.usc.edu). Amyloid PET scans were obtained with different tracers (*i.e.*, Pittsburgh compound B, Florbetapir, Florbetaben) depending on the ADNI study phase, and pre-processed by the ADNI PET imaging corelab (*i.e.*, ‘co-reg, avg, standardized image and voxel size’). In addition to imaging data, we used the cortical composite standardized uptake value ratios (SUVR) (Kinahan and Fletcher 2010; Landau et al. 2015) normalized by the whole cerebellum as a reference region, and added it as a standard amyloid measurement from PET in our model’s performance evaluation.

As some participants had several clinical visits, a total of 1,870 amyloid PET images were obtained. Among them individuals had AD (n=190), amnestic mild cognitive impairment (aMCI) due to AD (n=928), subjective memory complaints (SMC) (n=145) or were cognitively normal (CN) (n=607) based on the ADNI clinical classification (Table S2). The clinical classifications in ADNI are based on the Clinical Dementia Rating (CDR), the Mini-Mental State Examination (MMSE) and delayed recall of 1 paragraph from the Logical Memory (LM) II of the Wechsler Memory Scale-Revised (Petersen et al. 2010). In addition to clinical classifications, we used immediate and delayed recall measures from the LM Story A and also immediate (total words in trials 1-5) and delayed recall measures of the Rey Auditory Verbal Learning Test (AVLT): these tests were used as continuous measures of episodic memory.

### 2.2. Biological classification based on CSF measurements

The biological classes were defined based on earlier established cut-off values of amyloid-β 1-42 and phosphorylated tau CSF measurement (Shaw and Trojanowski 2017). In the first phases of ADNI, CSF quantification was implemented by INNO-BIA AlzBio3 immunoassay kit-based reagents, however, later the approach was replaced by fully automated Roche Elecsys. These methods have different scaling ranges, consequently, threshold values for biomarker aggregation also differ (amyloid-β: 192 pg/mL for INNO-BIA AlzBio3, 980 pg/mL for Roche Elecsys; tau: 93 pg/mL for INNO-BIA AlzBio3, 245 pg/mL for Roche Elecsys; phosphorylated tau (p-tau): 23 pg/mL for INNO-BIA AlzBio3, 21.8 pg/mL for Roche Elecsys) (Shaw et al. 2009; Shaw and Trojanowski 2017).

As the result we obtained four biological classes: participants with negative amyloid and tau proteins’ CSF measurements based on cut-off values (A-T-); amyloid negative, tau positive group (A-T+); amyloid positive, tau negative group (A+T-); amyloid and tau positive group (A+T+). We used only A and T in our biological classification, whereas the neurodegeneration (N) component was not included because it is not currently needed for biological classification of AD related pathology (Jack et al. 2018).

### 2.3. Fluid biomarker rescaling

To integrate values from the different CSF measurement methods, we trained linear regression and third-degree polynomial regression models to rescale AlzBio3 values to Roche Elecsys based on 1,072 samples with both values available (858 training samples, 214 test samples). The model which maximized the coefficient of determination (R^2^) and accuracy of positive and negative amyloid classification using cut-off values in the test dataset was selected for further analysis.

### 2.4. A deep neural network model to predict CSF values from PET scans

We developed a deep convolutional neural network model called “ArcheD” which utilizes residual connections (He et al. 2015) to predict logarithmic Aβ CSF values from amyloid PET scans. The residual connections allow for a deeper model to be trained which has been shown to improve performance (He et al. 2015). Our model architecture includes one initial convolution-pooling block followed by two residual-pooling blocks, two fully connected layers and a linear regression node (Fig. 1). Each residual block contains a sequence of convolutional, batch normalization and rectified linear activation function (ReLU) activation layers.

**Fig. 1.**
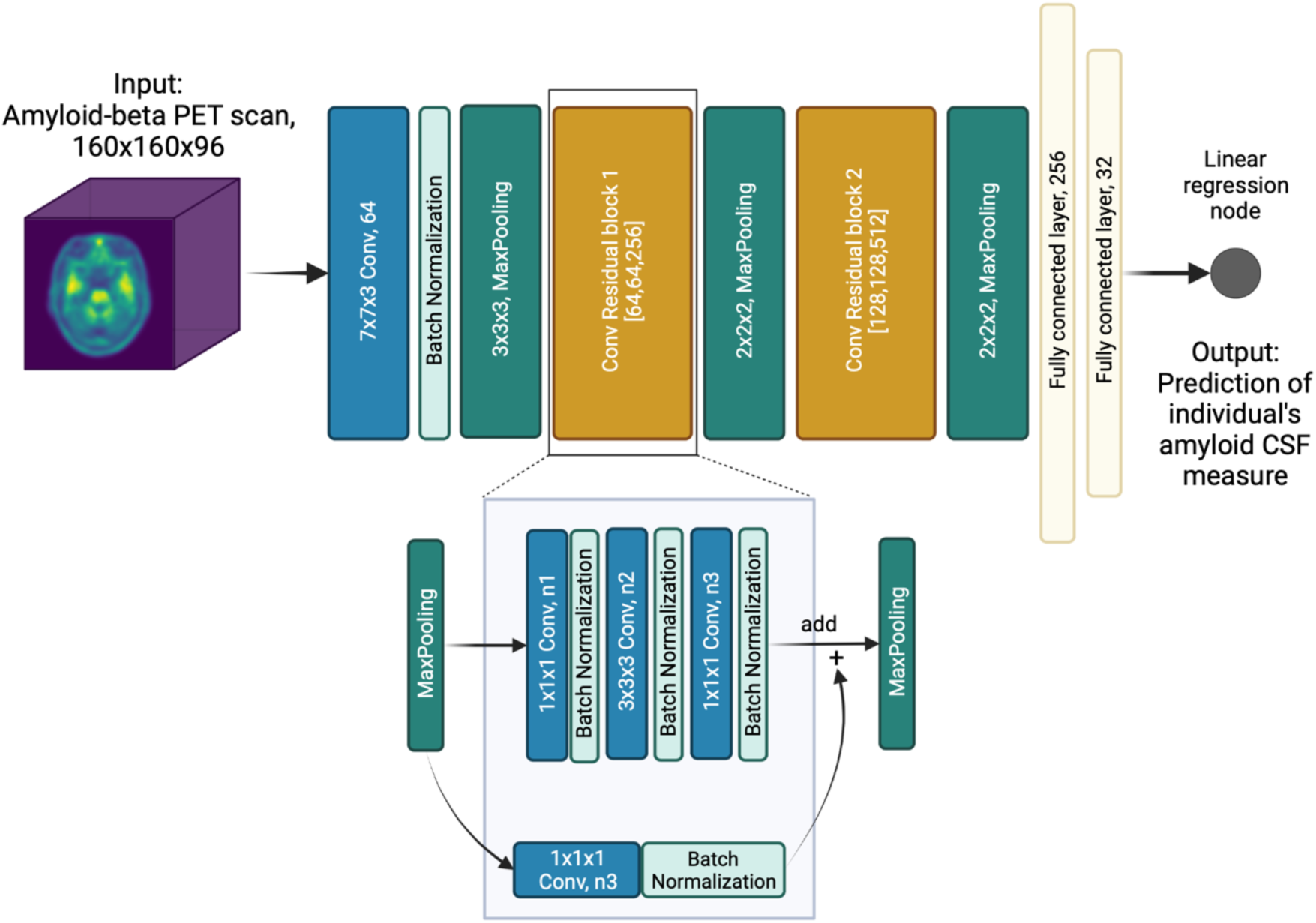
Schematic presentation of the ArcheD model architecture. PET – positron emission tomography; CSF – cerebrospinal fluid; n1, n2 and n3 – number of filters in convolution layers are specified in brackets for both convolutional residual blocks. Created with BioRender.com

ArcheD contains 35 layers and approximately 10.7 millions of parameters. To train the model, we used the Adam optimization algorithm (Kingma and Ba 2014) with an initial learning rate of 0.0001 to minimize the mean squared error (MSE) of observed and predicted CSF values. The model was trained for a maximum of 150 epochs with minibatch size of four, and stopped early, if loss in the validation dataset did not decrease for 15 epochs.

We used 60% of the dataset for training (n=1,197), 20% for validation (n=299) and 20% for testing the model (n=374). To increase the size of the training dataset with data augmentation, we either applied gaussian noise (σ=0,5,10,15,20,25%; six possible augmentations) or flipped the image by X or Y axes (two possible augmentations). To generate an augmented image from a training dataset image, one of the eight possible augmentations was selected with an equal probability. In the end, we obtained a total of 14,352 original and augmented brain scans constituting the augmented training data. To evaluate the robustness of ArcheD in data not used in training, we used a test dataset. ArcheD code is available at GITHUB (https://github.com/artagmaz/arched.git).

### 2.5. Guided backpropagation relevance maps

To explore which brain regions contribute the most to ArcheD’s predictions of amyloid CSF measurements from amyloid PET scans, we used the guided backpropagation technique (Springenberg et al. 2014). This approach quantifies how much model outputs change in response to perturbing model inputs. In PET imaging data, we used guided backpropagation to extract the contribution (relevance value) of each input voxel to CSF prediction. Guided backpropagation creates more specific relevance value maps compared to the classic backpropagation approach by replacing negative gradients with zero in ReLU activation layers during backward pass (Rieke et al. 2018) (Fig. 2.1).

**Fig. 2.**
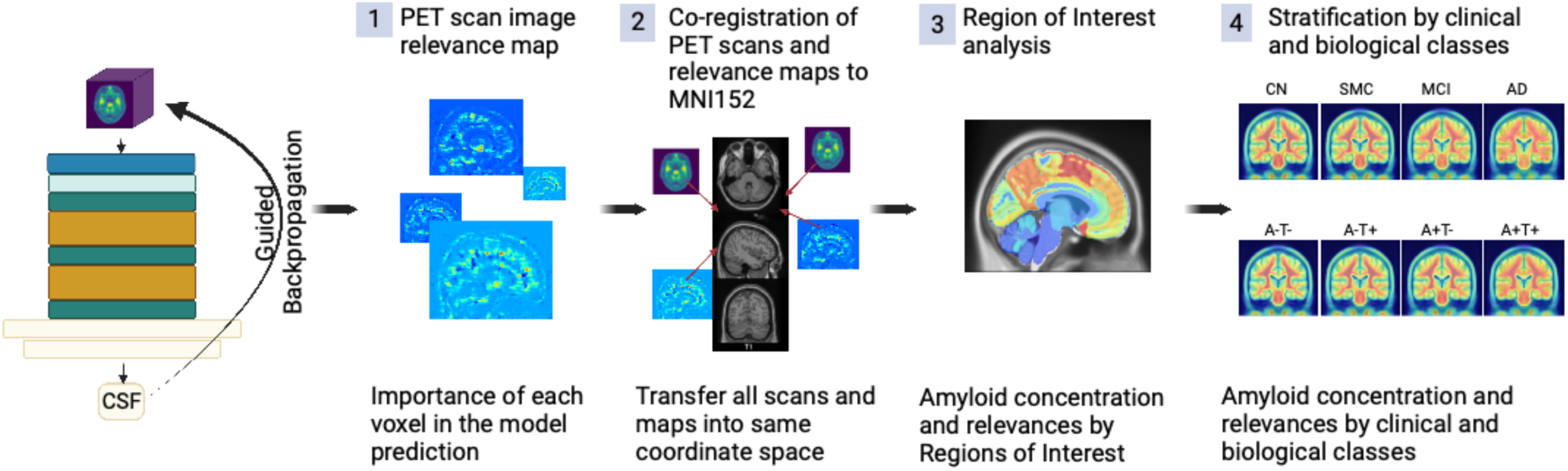
Workflow to process PET scans with ArcheD and guided backpropagation to identify characteristics distinguishing brain regions of interest and clinical and biological groups.

To compare the relevance maps between individuals and AD classes, we co-registered PET scans and corresponding relevance maps to individuals’ MRI and MNI152 templates with the AntsPy package (Avants, Tustison, and Song 2009) (Fig. 2.2). Since all PET images and relevance maps were described in the same coordinate space, we were able to create average PET and relevance maps for all samples and to separate AD classes. We used the Neuromorphometrics atlas to derive regions-of-interest (ROIs) (Bakker, Tiesinga, and Kötter 2015) (Table S1, Fig. 2.3) and compared their relevance to the model decision-making between classes (Fig. 2.4). The relevance value per ROI was normalized to the size of the region to evaluate how relevant the average voxel of each area is for CSF predictions.

### 2.6. Statistical analyses

We computed Pearson correlations with two-tailed p-values to quantify the strength of the associations between model prediction and CSF Aβ, cortical SUVR, and episodic memory measures. We also compared relevance values of brain areas between biological or clinical classes by bootstrapping with 95% confidence interval, Cohen’s *d* and Welch’s t-test. For all multiple comparisons, we computed false discovery rates (FDR) with the Benjamini-Hochberg method.

### 2.7. Programming environment

Tensorflow 2.4.1 and Python 3.9.7 were used to develop the machine learning models and perform computational analyses. The models were trained on NVIDIA Tesla V100 GPUs with 16 GB memory. For uploading, augmentation, registration and visualization of PET images we used nibabel, dltk and AntsPy python packages (Avants, Tustison, and Song 2009; Brett et al. 2023; Pawlowski et al. 2017). Functions for guided backpropagation gradient analysis were adapted from https://github.com/jrieke/cnn-interpretability. All R and Python scripts used in the study are provided at https://github.com/artagmaz/arched.git.

## 3. Results

The mean age of the participants was 73.6 years (SD =7.37) and 49% were women. Detailed descriptive statistics of study participants are presented in Table S2.

### 3.1. Model training

#### 3.1.1. Fluid biomarker rescaling

We first transformed AlzBio3 CSF values to conform to the range of values present in Roche Elecsys CSF measurements (Shaw and Trojanowski 2017). Both regression models performed similarly on a held-out portion of the data (linear regression: R^2^test=68.9%, 91.5% accuracy; 3rd degree polynomial regression: R^2^test=69.4%, 91.5% accuracy) (Table S3, Figure S1). Based on these metrics and visual inspection of the regression models (Fig. 3.A), we decided to use a 3rd degree polynomial regression model to predict Roche Elecsys amyloid measurements for the remaining 1,102 samples.

**Fig. 3.**
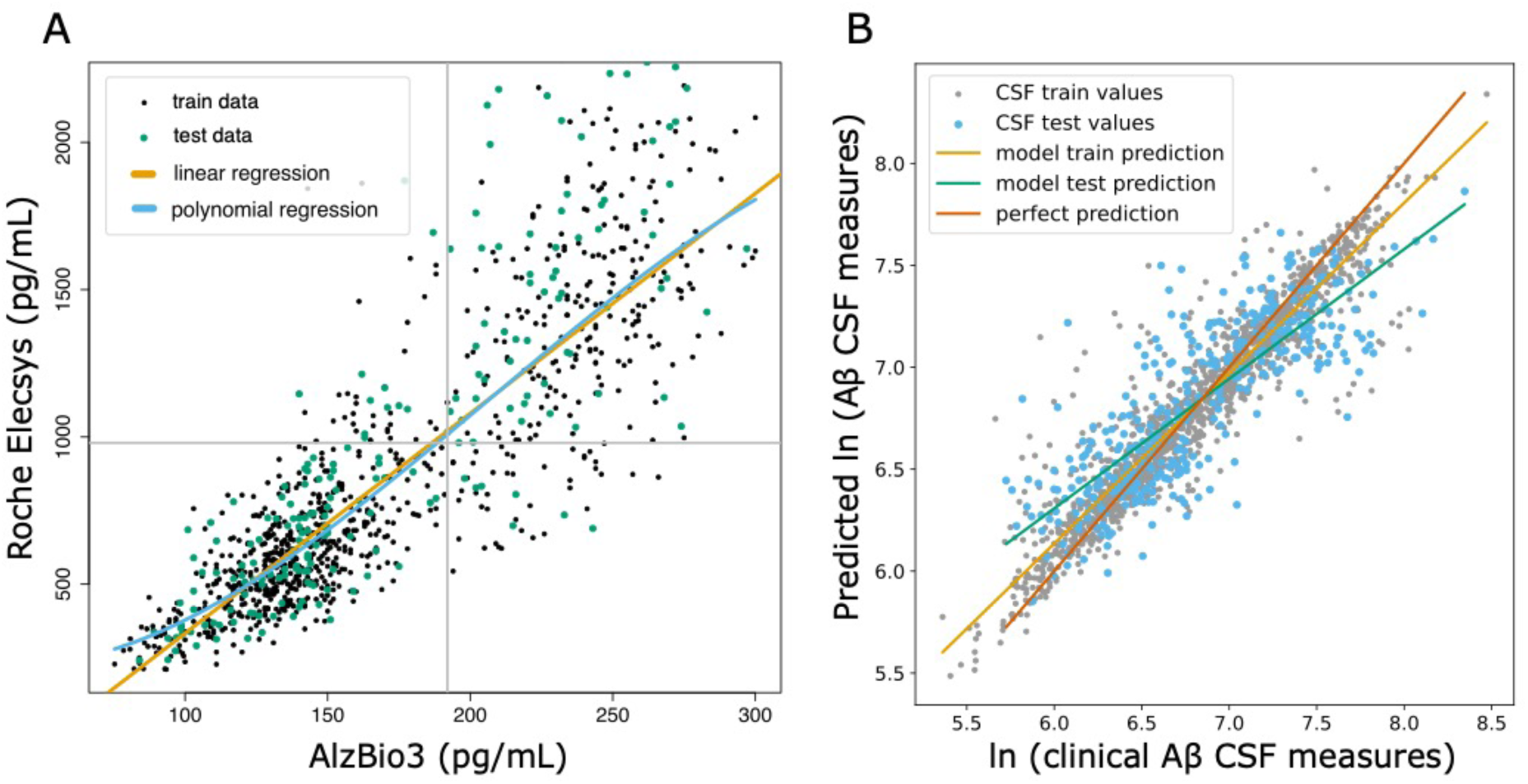
**(A)** Linear (orange line) and 3rd degree polynomial (blue line) regression for fluid amyloid rescaling from INNO-BIA AlzBio3 system to Roche Elecsys. Gray lines present cut-off values for Aβ CSF measurement. **(B)** Comparison of clinically measured Aβ CSF and values predicted by the ArcheD model. CSF values given as natural logarithms. Lines illustrate the linear relationship between clinical Aβ CSF measurement and prediction of our model: green ‒ linear relation between real and predicted values of the test dataset; orange ‒ linear relation between real and predicted values of the training dataset. Red diagonal line shows the ideal one-to-one correspondence between observed and predicted values.

#### 3.1.2. Training the ArcheD model and overall predictive performance

ArcheD was trained on 14,352 augmented PET scan images to predict logarithmic CSF values. The model achieving the best performance in the validation set was obtained after the eighth epoch of training (mean squared error, MSE=0.12) (Figure S1). Finally, ArcheD explained 66% of the variance (R^2^) in the test dataset not used in training (Fig. 3.B).

### 3.2. Associations of DL with SUVR and episodic memory measurements

The ArcheD Aβ CSF prediction was correlated with biological and cognitive AD markers (Table 1, Figure S2). A significant association was observed between predicted and clinically measured CSF Aβ, both for all samples and all clinical classes (*r*>0.89, *q*<0.01). SUVR strongly correlated with all sample subsets’ Aβ CSF (*r*<-0.53, *q*<0.01) except for AD samples (*r*av45=-0.42, *q*<0.05; *r*fbb=-0.08, *q*=0.76). In the case of episodic memory test results, CSF predictions were generally weaker and mostly non-significant in the AD (*n*=190) and SMC (*n*=145) groups compared to MCI (*n*=928) and CN (*n*=607) groups. There was a significant positive correlation in AD samples between predicted Aβ CSF and AVLT immediate recall (*r=*0.36; *q<*0.01), whereas in those with SMC predicted CSF measurements showed no significant correlations with any of the episodic memory measures (Table 1). All memory test scores for MCI and CN groups were significantly correlated with predicted amyloid CSF measurement (*r>*0.16, *q<*0.01) (Table 1).

**Table 1.**
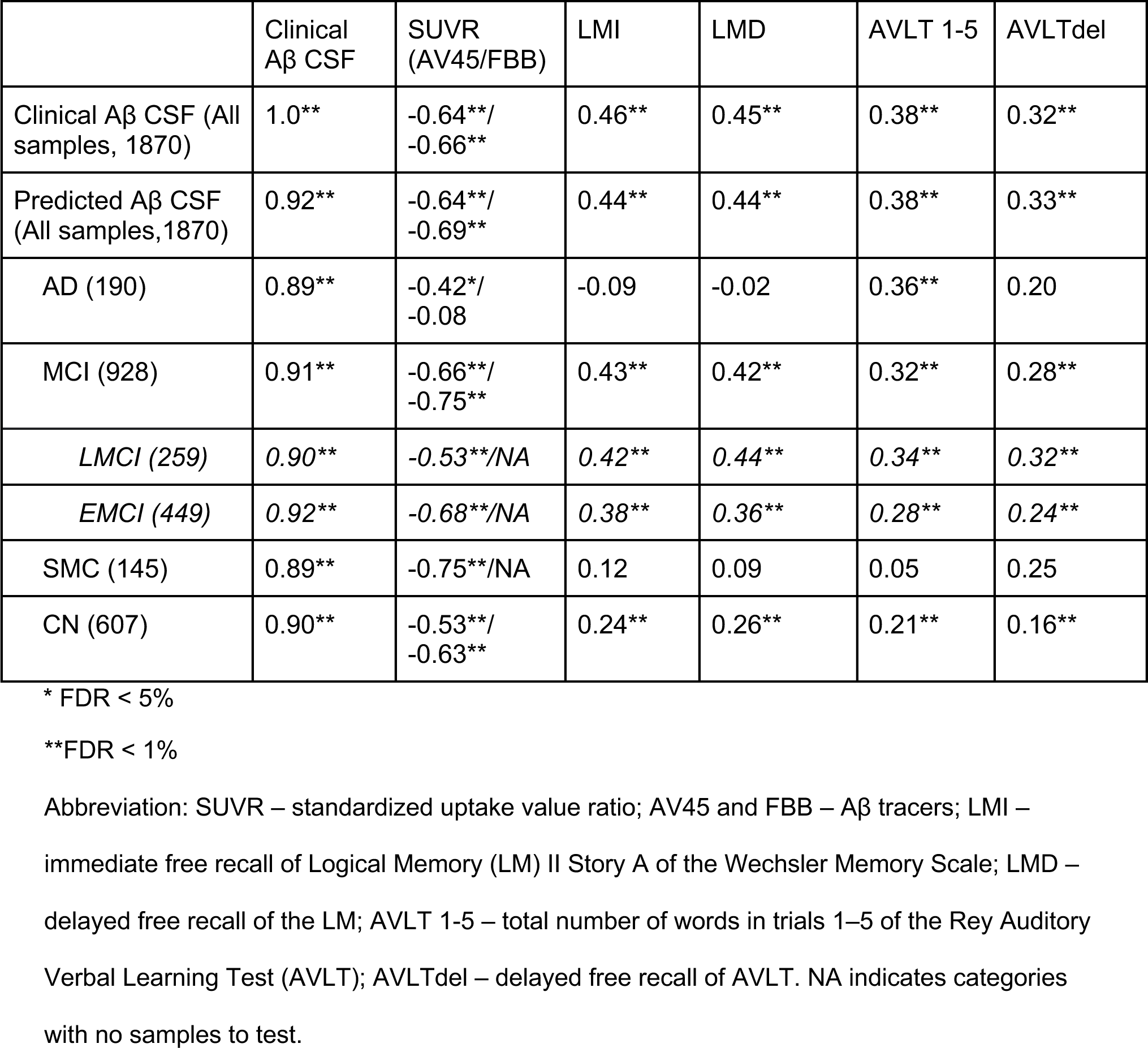
Pearson correlations between Aβ CSF values predicted by model and clinical, imaging and memory markers of AD.

### 3.3. Relevance of brain areas for model decision making

To understand which brain regions contributed the most to ArcheD predictions, we compared the relevance of these regions with guided backpropagation. We found that the areas which contributed the most were cerebral (mean relevance 0.251, 95% CI [0.247, 0.255]) and cerebellum white matter (0.207, 95% CI [0.205;0.208]) followed by brain stem, subcortical areas, gray matter regions at the lobar level and cerebellum gray matter, whereas limbic lobe, basal forebrain, ventricles and optic chiasm were substantially less important for the model (Table S4, Fig. 4.A-B).

**Fig 4.**
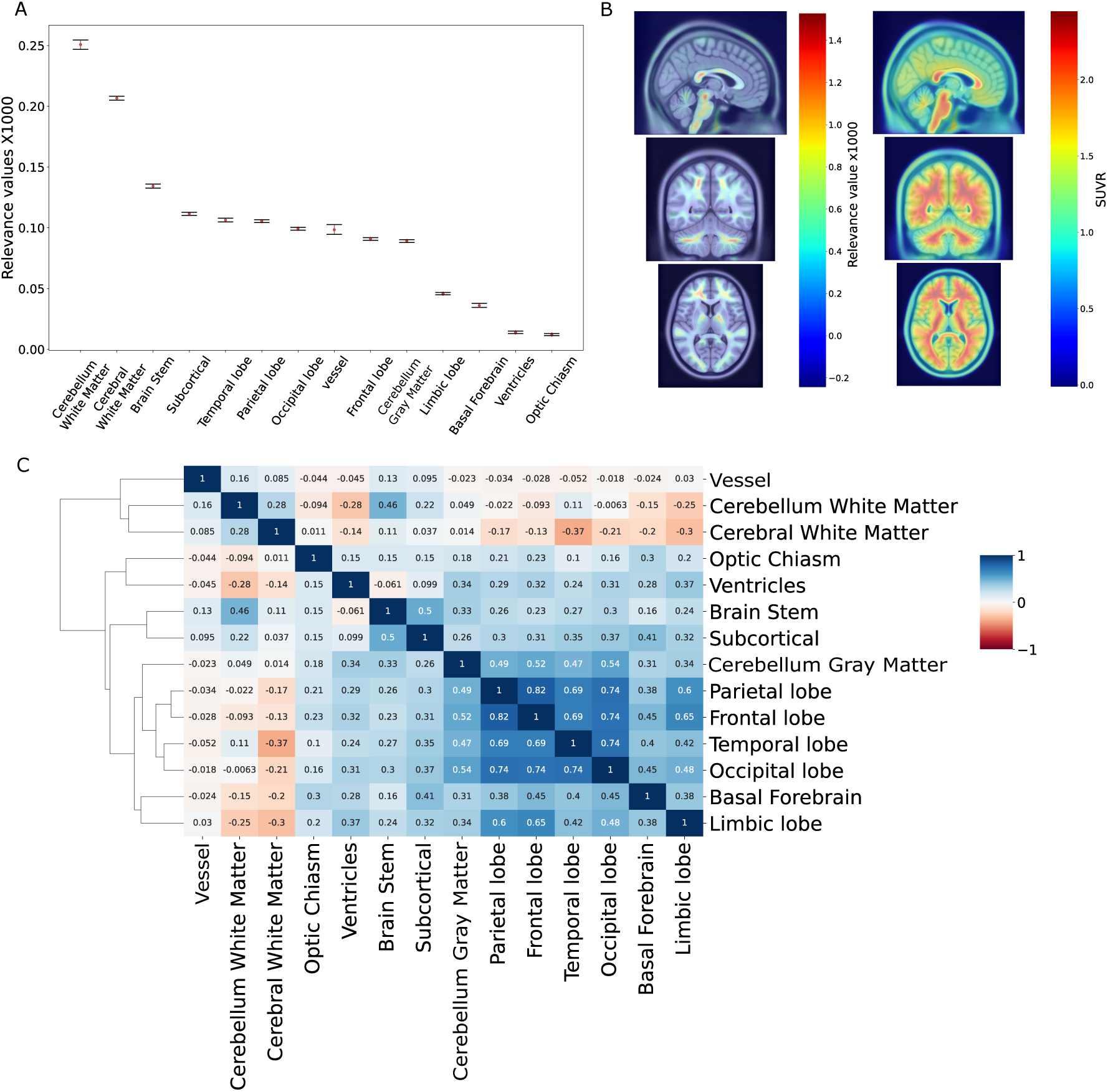
**(A)** Mean relevance values for brain regions and bootstrapped 95% confidence intervals. Values were multiplied by 1000. **(B)** Average relevance value and amyloid concentration (SUVR) for the dataset on PET scan. **(C)** Heatmap of Pearson correlations for relevance values of all brain regions.

We then examined the similarity of ROIs based on their relevance profiles in the ADNI cohort (Fig. 4.C). As expected, cerebral and cerebellum white matter, and vessels clustered together. The main cerebral lobes, *i.e.*, frontal, temporal, parietal and occipital lobe were clustered together, as were the brainstem and subcortical regions. The remaining regions formed two clusters containing optic chiasm and ventricles, and basal forebrain and limbic lobe (Fig. 4.C).

### 3.4. Region specific contributions to prediction of CSF in clinical and biological sub-classes

The between classes analysis showed that there were brain areas that contributed at the same level for all clinical or biological classes (optic chiasm, ventricles, and vessels), and areas that were significantly different between classes by relevance value (Figure S3, Fig. 5, Table S4). Significant differences in relevance values between AD/MCI and CN groups were observed in eight ROIs (*q*<0.01) (Table S5). Cerebral white matter contributed more to Aβ CSF value predictions in the CN group (AD vs CN Cohen’s *d*=-0.976, A+T+ vs A-T-*d*=-1.138; *q*<0.01), whereas other regions (cortical lobes, limbic lobe, basal forebrain and subcortical areas) were more important for prediction in the AD and MCI groups (Table S5). Brainstem relevance values differentiated only between MCI and CN groups (Fig. 5, Table S5). However, there were no significant differences between SMC and CN groups (Fig. 5, Table S5).

**Fig. 5.**
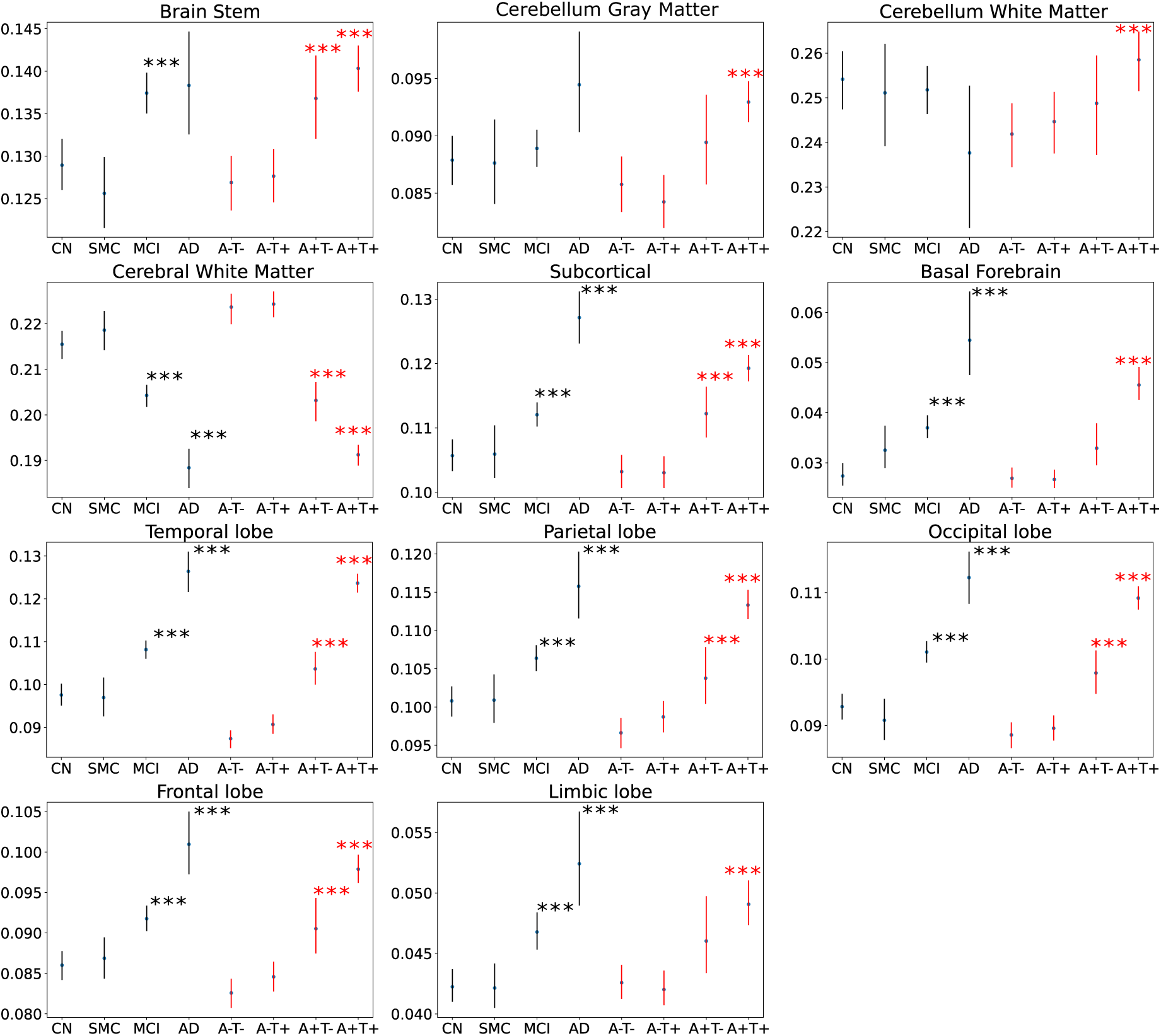
Comparison of mean relevance values per brain region within clinical or biological classifications. Cognitive normal (CN) group was used as control for clinical classification. A-T- was used as control for biological classification. *** ‒ *q*<0.01.

Most of the relevance patterns observed for biological classes were concordant with those seen in clinical classes. In all ROIs, except vessels, ventricles and optic chiasm, a marked difference in relevance values was visible between A+T+ and A-T- classes (Fig. 5, Table S5). In addition, parietal, occipital, frontal, temporal lobes, brain stem, cerebral white matter and subcortical areas differentiated A+T- from the A-T- (*q*<0.01) (Fig. 5, Table S5).

### 3.5. Contribution of gray matter regions to Aβ CSF prediction in clinical and biological classes

Taking in account the high amount of unspecified binding of the amyloid tracers in cerebral and cerebellum white matter, we conducted a closer investigation focusing only on cortical gray matter (GM) regions (Fig. 4.B) (Klunk et al. 2004).

At the lobar level, temporal (mean relevance 0.106, 95% CI [0.105, 0.108]) and parietal (0.105, 95% CI [0.104, 0.107]) lobes contributed the most to the model prediction. In contrast, occipital (0.098, 95% CI [0.098, 0.100]) and frontal lobes (0.091, 95% CI [0.089, 0.92]) had significantly lower relevance values than temporal and parietal lobes (Fig. 6.A). However, when looking at absolute levels of PET amyloid binding as expressed by SUVR, parietal (mean SUVR 1.449, 95% CI [1.432, 1.466]), occipital (mean 1.440, 95% CI [1.426, 1.455]) and frontal (1.435, 95% CI [1.419, 1.452]) lobes showed higher levels of amyloid aggregation compared to temporal lobe (1.359, 95% CI [1.344, 1.375]) (Fig. 6.A).

**Fig. 6.**
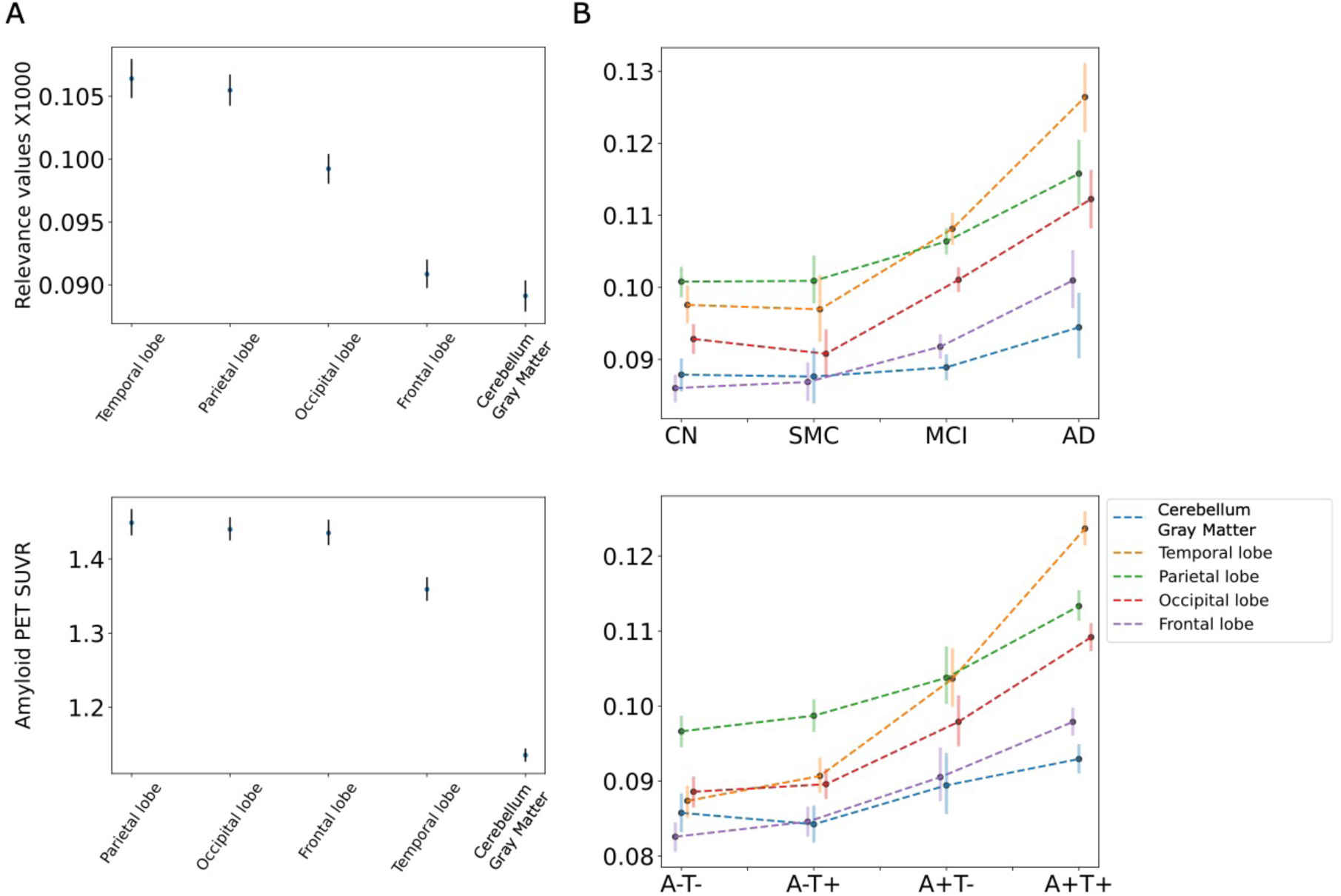
**(A)** Mean relevance values (top) and amyloid concentration (bottom) for gray matter regions and bootstrapped 95% confidence intervals. **(B)** Relevance level of gray matter regions on different clinical (top) and biological classes (bottom) of AD.

When comparing relevance values differences between biological or clinical classes, we observed that the most contributing gray matter region differed along the AD continuum (Fig. 6.B, Table S6). In the CN and SMC groups (based on the clinical classification), the parietal lobe (CN mean 0.101, 95% CI [0.099, 0.103]; SMC 0.101, 95% CI [0.098, 0.104]) was the most significantly contributing region, followed by the temporal, occipital lobes, cerebellum gray matter and frontal lobe (Fig. 6.B, Table S6). In MCI and AD groups, the relevance values of all regions were higher than in the CN group, and the temporal lobe contributed more than the other lobes (MCI temporal 0.108, 95% CI [0.106, 0.110]; AD temporal 0.126, 95% CI [0.122, 0.131]).

For biological classification, we observed a similar pattern in terms of the early versus late stage of the AD continuum. In the A-T- group, the parietal lobe (0.097, 95% CI [0.095, 0.098]) contributed most significantly to the model decision-making, followed by the occipital, temporal lobes, cerebellum gray matter and frontal lobe (Fig. 6.B, Table S6). In the A-T+ and A+T- groups, the temporal lobe was more relevant than occipital area (A-T+ temporal 0.091, 95% CI [0.089, 0.093]; A+T- temporal 0.104, 95% CI [0.100, 0.108]), and dominated all other gray matter regions in A+T+ class (0.124, 95% CI [0.122, 0.126]) (Fig. 6.B, Table S6).

## 4. Discussion

We developed a novel deep neural network model (ArcheD) for prediction of CSF amyloid biomarker concentration directly from amyloid PET images independent of the amyloid PET tracer. Trained on ADNI PET images and associated Aβ CSF measurements, ArcheD was able to explain 66% of variance in CSF levels in a test dataset withheld from training. We thus envision using ArcheD to complement the information obtained in a Aβ PET scan by providing an estimate of the Aβ CSF level without a lumbar puncture.

CSF Aβ predicted by our model correlated significantly with cortical amyloid PET SUVR and episodic memory performance. These correlations were similar to correlations between biological and clinical measures (amyloid PET SUVR and episodic memory measurements) that are used to determine the AD hallmarks of amyloid positivity and episodic memory impairment (Hake et al. 2015; Niemantsverdriet et al. 2017). The absence of association in AD and SMC groups with FBB SUVR and episodic memory performance could be explained by the smaller number of participants in these classes compared to group sizes of CN and MCI. In addition, the AD group included individuals with severe AD, resulting in little variation in amyloid levels and episodic memory performance; amyloid levels plateau in the course of AD and there is also little variation in delayed recall measures in those with severe AD (Klunk et al. 2004; Thal et al. 2002).

To understand which brain regions were the most influential in predicting CSF Aβ levels, we investigated the relevance of average voxel per brain region and compared ROIs between each other, and furthermore, performed these comparisons separately in clinical and biological sub-classes. Cerebellum and cerebral white matter, followed by subcortical area and brainstem were identified as the most influential brain regions for model predictions. These regions have unspecified binding of amyloid tracers (Klunk et al. 2004; Matsubara et al. 2016) and can be used as visual control regions to test the tracers’ delivery to the brain. They are also not affected by amyloid plaques until the very late AD stage (Klunk et al. 2004; Thal et al. 2002) and can be used as possible reference regions for calculating SUVR (Heeman et al. 2020), even if cerebellum gray matter is the gold standard reference region. It is likely that our ArcheD model learnt to use these regions as a reference point for CSF prediction and combined them with cortical regions to calibrate amyloid CSF values. The areas contributing the least to predictions were limbic lobe, basal forebrain, ventricles and optic chiasm. These ROIs are known to be less involved in the amyloid accumulation process during the AD development (Hampel, Hardy, et al. 2021).

Furthermore, we explored if the same or different regions contributed to CSF prediction in different clinical and biological classes. Relevance of cerebral cortex (including limbic lobe), subcortical, brain stem, basal forebrain, cerebellum white and gray matter regions was significantly greater in those with AD compared to those without AD or in those who were amyloid positive compared to those who were amyloid negative. Cerebral white matter contributed more to CSF prediction in individuals without cognitive impairment (CN, SMC) or in individuals who were amyloid negative (A-T-, A-T+). According to criterias for amyloid PET scans usage, amyloid-positive scans differentiate from amyloid-negative scans based on gray matter cortical uptake that is above the level of nonspecific binding in white matter (Johnson et al. 2013; Wolk et al. 2012; Richards and Sabbagh 2014). This corresponds to our finding, indicating that cerebral white matter plays a more significant role in CSF prediction for amyloid-negative groups, while cortical regions are more relevant to amyloid positive individuals. We hypothesized that ArcheD might not only use cerebral white matter as a reference region but also can focus on both white and gray matter amyloid binding levels together.

Focusing on cortical GM, we found considerable differences between lobes’ relevance values and amyloid deposition level. Temporal and parietal areas contributed the most to the model prediction, whereas occipital and frontal lobes’ relevance values were noticeably less. Temporal lobe has been reported to substantially contribute to model decision-making also in other studies of AD classification based on MRI scans (Dyrba et al. 2021; Rieke et al. 2018). However, on PET scans, parietal, occipital and frontal lobes have higher levels of amyloid aggregation, while temporal lobe has significantly less amyloid. According to the Aβ pathology studies, parietal and frontal lobes are regions with early amyloid accumulation which can explain the high biomarker concentration in those regions in our results (Insel et al. 2020; Mattsson et al. 2019).

Despite temporal lobe being the least amyloid enriched lobe in PET scans, ArcheD highlighted this lobe as the most important cortical region for the amyloid CSF prediction. When we looked at the level of cortical lobes’ relevance between different classes, we noticed that temporal lobe’s relevance is more important than other lobes only in MCI, AD and A+T+ groups which are indicating later stages in the AD continuum compared to SMC, CN and A-negative groups. In more early stages of the disease development the parietal lobe is the most significant contributor, which is compatible with earlier studies (Insel et al. 2020; Mattsson et al. 2019).

This study presented a novel neural network model that analyzes amyloid PET scans, independent of the amyloid tracer used, and predicts what the amyloid beta level would be in CSF. The model learned to focus on specific brain regions depending on the AD stage, defined by the biological or clinical classification. These regions aligned well with previous findings on AD progression.

ArcheD has the potential to be used in clinical practice in the future. It can reduce the amount of work for clinicians and potentially prevent patients from lumbar puncture. ArcheD is also straightforward to use, as the PET scan image is the only input to the method. However, our research has some limitations that can be improved in future studies. We had only a limited dataset of PET scans with available CSF measurements, especially of AD individuals, to train the ArcheD model. Larger training datasets would likely yield better predictive performance. Another limiting factor is the strict requirement for input PET images to have dimensions of 160×160×96 voxels. Therefore, users may need to perform preprocessing, such as co-registration, averaging, standardization, and voxel size adjustment, similar to ADNI PET imaging corelab, on their own. Our model and findings also need to be validated on external datasets.

## 5. Conclusion

In conclusion, ArcheD was able to predict amyloid CSF values directly from amyloid PET scans in the AD continuum (including cognitively unimpaired individuals with preclinical AD). Predicted CSF values correlated significantly with CSF amyloid, cortical amyloid SUVR and episodic memory performance. It was also comparable with the correlations between CSF amyloid and SUVR as well as episodic memory. Analyzing the sensitivity of CSF predictions to input PET images by brain region, cerebellum white matter, cerebral white matter, brainstem and subcortical areas were found to contribute the most to the model prediction and may have been used by ArcheD as reference points for prediction. We compared contribution of brain regions to predictions within clinical and biological classification, and found that the cerebral cortex, subcortical areas, brain stem, cerebellum white and gray matter, and basal forebrain influenced predictions especially in the MCI, AD, A+T- and A+T+ classes, while cerebral white matter contributed more in clinically normal and early-stage groups. Taking a closer look at the cortical regions, we found that the focus of the model was in the parietal lobe in preclinical AD stage (CN, SMC) and in those who were amyloid or tau negative (A-T-, A-T+, A+T-) whereas temporal lobe was most implicated in clinical groups with memory impairment (MCI, AD) and in those who were both amyloid and tau positive (A+T+). Our model can serve in clinical practice for determining an Aβ CSF state and improving AD early detection. However, further studies are needed to validate and tune the model for clinical use.

## Acknowledgements

The authors wish to acknowledge CSC – IT Center for Science, Finland, for generous computational resources.

Supported by the Doctoral Programme in Population Health of the University of Helsinki (AT), and the Academy of Finland (grants 314639, 320109 and 345988 to EV, and 322675 and 328890 to EP).

Data collection and sharing for this project was funded by the Alzheimer’s Disease Neuroimaging Initiative (ADNI) (National Institutes of Health Grant U01 AG024904) and DOD ADNI (Department of Defense award number W81XWH-12-2-0012). ADNI is funded by the National Institute on Aging, the National Institute of Biomedical Imaging and Bioengineering, and through generous contributions from the following: AbbVie, Alzheimer’s Association; Alzheimer’s Drug Discovery Foundation; Araclon Biotech; BioClinica, Inc.; Biogen; Bristol-Myers Squibb Company; CereSpir, Inc.; Cogstate; Eisai Inc.; Elan Pharmaceuticals, Inc.; Eli Lilly and Company; EuroImmun; F. Hoffmann-La Roche Ltd and its affiliated company Genentech, Inc.; Fujirebio; GE Healthcare; IXICO Ltd.; Janssen Alzheimer Immunotherapy Research & Development, LLC.; Johnson & Johnson Pharmaceutical Research & Development LLC.; Lumosity; Lundbeck; Merck & Co., Inc.; Meso Scale Diagnostics, LLC.; NeuroRx Research; Neurotrack Technologies; Novartis Pharmaceuticals Corporation; Pfizer Inc.; Piramal Imaging; Servier; Takeda Pharmaceutical Company; and Transition Therapeutics. The Canadian Institutes of Health Research is providing funds to support ADNI clinical sites in Canada. Private sector contributions are facilitated by the Foundation for the National Institutes of Health (www.fnih.org). The grantee organization is the Northern California Institute for Research and Education, and the study is coordinated by the Alzheimer’s Therapeutic Research Institute at the University of Southern California. ADNI data are disseminated by the Laboratory for Neuro Imaging at the University of Southern California.

## References

Aizenstein, Howard Jay, Robert D. Nebes, Judith A. Saxton, Julie C. Price, Chester A. Mathis, Nicholas D. Tsopelas, Scott K. Ziolko, et al. 2008. “Frequent Amyloid Deposition without Significant Cognitive Impairment among the Elderly.” Archives of Neurology 65 (11): 1509–17.

Avants, Brian, Nicholas J. Tustison, and Gang Song. 2009. “Advanced Normalization Tools: V1.0.” The Insight Journal, July. 10.54294/uvnhin.

Bakker, Rembrandt, Paul Tiesinga, and Rolf Kötter. 2015. “The Scalable Brain Atlas: Instant Web-Based Access to Public Brain Atlases and Related Content.” Neuroinformatics 13 (3): 353–66.

Brett, Matthew, Christopher J. Markiewicz, Michael Hanke, Marc-Alexandre Côté, Ben Cipollini, Paul McCarthy, Dorota Jarecka, et al. 2023. “Nipy/nibabel: 5.1.0,” April. 10.5281/zenodo.7795644.

Choi, Hongyoon, Yu Kyeong Kim, Eun Jin Yoon, Jee-Young Lee, Dong Soo Lee, and Alzheimer’s Disease Neuroimaging Initiative. 2020. “Cognitive Signature of Brain FDG PET Based on Deep Learning: Domain Transfer from Alzheimer’s Disease to Parkinson’s Disease.” European Journal of Nuclear Medicine and Molecular Imaging 47 (2): 403–12.

Ding, Yiming, Jae Ho Sohn, Michael G. Kawczynski, Hari Trivedi, Roy Harnish, Nathaniel W. Jenkins, Dmytro Lituiev, et al. 2019. “A Deep Learning Model to Predict a Diagnosis of Alzheimer Disease by Using F-FDG PET of the Brain.” Radiology 290 (2): 456–64.

Dyrba, Martin, Moritz Hanzig, Slawek Altenstein, Sebastian Bader, Tommaso Ballarini, Frederic Brosseron, Katharina Buerger, et al. 2021. “Improving 3D Convolutional Neural Network Comprehensibility via Interactive Visualization of Relevance Maps: Evaluation in Alzheimer’s Disease.” Alzheimer’s Research & Therapy 13 (1): 1–18.

Erickson, Kirk I., Shannon D. Donofry, Kelsey R. Sewell, Belinda M. Brown, and Chelsea M. Stillman. 2022. “Cognitive Aging and the Promise of Physical Activity.” Annual Review of Clinical Psychology 18 (May): 417–42.

Hake, Ann, Paula T. Trzepacz, Shufang Wang, Peng Yu, Michael Case, Helen Hochstetler, Michael M. Witte, Elisabeth K. Degenhardt, Robert A. Dean, and Alzheimer’s Disease Neuroimaging Initiative. 2015. “Florbetapir Positron Emission Tomography and Cerebrospinal Fluid Biomarkers.” Alzheimer’s & Dementia 11 (8): 986–93.

Hampel, Harald, Rhoda Au, Soeren Mattke, Wiesje M. van der Flier, Paul Aisen, Liana Apostolova, Christopher Chen, et al. 2022. “Designing the next-Generation Clinical Care Pathway for Alzheimer’s Disease.” Nature Aging 2 (8): 692–703.

Hampel, Harald, Jeffrey Cummings, Kaj Blennow, Peng Gao, Clifford R. Jack Jr, and Andrea Vergallo. 2021. “Developing the ATX(N) Classification for Use across the Alzheimer Disease Continuum.” Nature Reviews. Neurology 17 (9): 580–89.

Hampel, Harald, John Hardy, Kaj Blennow, Christopher Chen, George Perry, Seung Hyun Kim, Victor L. Villemagne, et al. 2021. “The Amyloid-β Pathway in Alzheimer’s Disease.” Molecular Psychiatry 26 (10): 5481–5503.

Heeman, Fiona, Janine Hendriks, Isadora Lopes Alves, Rik Ossenkoppele, Nelleke Tolboom, Bart N. M. van Berckel, Adriaan A. Lammertsma, Maqsood Yaqub, and on behalf of the AMYPAD Consortium. 2020. “[11C]PIB Amyloid Quantification: Effect of Reference Region Selection.” EJNMMI Research 10 (1). 10.1186/s13550-020-00714-1.

He, Kaiming, Xiangyu Zhang, Shaoqing Ren, and Jian Sun. 2015. “Deep Residual Learning for Image Recognition,” December. 10.48550/arXiv.1512.03385.

Insel, Philip S., Elizabeth C. Mormino, Paul S. Aisen, Wesley K. Thompson, and Michael C. Donohue. 2020. “Neuroanatomical Spread of Amyloid β and Tau in Alzheimer’s Disease: Implications for Primary Prevention.” Brain Communications 2 (1). 10.1093/braincomms/fcaa007.

Jack, Clifford R., Jr, Marilyn S. Albert, David S. Knopman, Guy M. McKhann, Reisa A. Sperling, Maria C. Carrillo, Bill Thies, and Creighton H. Phelps. 2011. “Introduction to the Recommendations from the National Institute on Aging-Alzheimer’s Association Workgroups on Diagnostic Guidelines for Alzheimer’s Disease.” Alzheimer’s & Dementia: The Journal of the Alzheimer’s Association 7 (3): 257–62.

Jack, Clifford R., Jr, David A. Bennett, Kaj Blennow, Maria C. Carrillo, Billy Dunn, Samantha Budd Haeberlein, David M. Holtzman, et al. 2018. “NIA-AA Research Framework: Toward a Biological Definition of Alzheimer’s Disease.” Alzheimer’s & Dementia: The Journal of the Alzheimer’s Association 14 (4): 535–62.

Jack, Clifford R., Jr, Heather J. Wiste, Timothy G. Lesnick, Stephen D. Weigand, David S. Knopman, Prashanthi Vemuri, Vernon S. Pankratz, et al. 2013. “Brain β-Amyloid Load Approaches a Plateau.” Neurology 80 (10): 890–96.

Jack, Clifford R., David S. Knopman, William J. Jagust, Leslie M. Shaw, Paul S. Aisen, Michael W. Weiner, Ronald C. Petersen, and John Q. Trojanowski. 2010. “Hypothetical Model of Dynamic Biomarkers of the Alzheimer’s Pathological Cascade.” The Lancet Neurology 9 (1): 119–28.

Johnson, Keith A., Satoshi Minoshima, Nicolaas I. Bohnen, Kevin J. Donohoe, Norman L. Foster, Peter Herscovitch, Jason H. Karlawish, et al. 2013. “Appropriate Use Criteria for Amyloid PET: A Report of the Amyloid Imaging Task Force, the Society of Nuclear Medicine and Molecular Imaging, and the Alzheimer’s Association.” Alzheimer’s & Dementia: The Journal of the Alzheimer’s Association 9 (1): e – 1–16.

Jo, Taeho, Kwangsik Nho, Shannon L. Risacher, Andrew J. Saykin, and Alzheimer’s Neuroimaging Initiative. 2020. “Deep Learning Detection of Informative Features in Tau PET for Alzheimer’s Disease Classification.” BMC Bioinformatics 21 (Suppl 21): 496.

Jo, Taeho, Kwangsik Nho, and Andrew J. Saykin. 2019. “Deep Learning in Alzheimer’s Disease: Diagnostic Classification and Prognostic Prediction Using Neuroimaging Data.” Frontiers in Aging Neuroscience 11 (August): 220.

Kim, Ji-Young, Hoon Young Suh, Hyun Gee Ryoo, Dongkyu Oh, Hongyoon Choi, Jin Chul Paeng, Gi Jeong Cheon, Keon Wook Kang, Dong Soo Lee, and Alzheimer’s Disease Neuroimaging Initiative. 2019. “Amyloid PET Quantification Via End-to-End Training of a Deep Learning.” Nuclear Medicine and Molecular Imaging 53 (5): 340–48.

Kinahan, Paul E., and James W. Fletcher. 2010. “Positron Emission Tomography-Computed Tomography Standardized Uptake Values in Clinical Practice and Assessing Response to Therapy.” Seminars in Ultrasound, CT, and MR 31 (6): 496–505.

Kingma, Diederik P., and Jimmy Ba. 2014. “Adam: A Method for Stochastic Optimization,” December. 10.48550/arXiv.1412.6980.

Klunk, William E., Henry Engler, Agneta Nordberg, Yanming Wang, Gunnar Blomqvist, Daniel P. Holt, Mats Bergström, et al. 2004. “Imaging Brain Amyloid in Alzheimer’s Disease with Pittsburgh Compound-B.” Annals of Neurology 55 (3): 306–19.

Knopman, D. S., J. E. Parisi, A. Salviati, M. Floriach-Robert, B. F. Boeve, R. J. Ivnik, G. E. Smith, et al. 2003. “Neuropathology of Cognitively Normal Elderly.” Journal of Neuropathology & Experimental Neurology 62 (11): 1087–95.

Landau, Susan M., Allison Fero, Suzanne L. Baker, Robert Koeppe, Mark Mintun, Kewei Chen, Eric M. Reiman, and William J. Jagust. 2015. “Measurement of Longitudinal β-Amyloid Change with 18F-Florbetapir PET and Standardized Uptake Value Ratios.” *Journal of Nuclear Medicine: Official Publication*, Society of Nuclear Medicine 56 (4): 567–74.

Lin, Eugene, Chieh-Hsin Lin, and Hsien-Yuan Lane. 2021. “Deep Learning with Neuroimaging and Genomics in Alzheimer’s Disease.” International Journal of Molecular Sciences 22 (15). 10.3390/ijms22157911.

Matsubara, Keisuke, Masanobu Ibaraki, Hitoshi Shimada, Yoko Ikoma, Tetsuya Suhara, Toshibumi Kinoshita, and Hiroshi Ito. 2016. “Impact of Spillover from White Matter by Partial Volume Effect on Quantification of Amyloid Deposition with [11C]PiB PET.” NeuroImage 143 (December): 316–24.

Mattsson, Niklas, Sebastian Palmqvist, Erik Stomrud, Jacob Vogel, and Oskar Hansson. 2019. “Staging β-Amyloid Pathology With Amyloid Positron Emission Tomography.” JAMA Neurology 76 (11): 1319–29.

McKhann, G., D. Drachman, M. Folstein, R. Katzman, D. Price, and E. M. Stadlan. 1984. “Clinical Diagnosis of Alzheimer’s Disease: Report of the NINCDS-ADRDA Work Group under the Auspices of Department of Health and Human Services Task Force on Alzheimer’s Disease.” Neurology 34 (7): 939–44.

Nelson, Peter T., Elizabeth Head, Frederick A. Schmitt, Paulina R. Davis, Janna H. Neltner, Gregory A. Jicha, Erin L. Abner, et al. 2011. “Alzheimer’s Disease Is Not ‘brain Aging’: Neuropathological, Genetic, and Epidemiological Human Studies.” Acta Neuropathologica 121 (5): 571–87.

Niemantsverdriet, Ellis, Julie Ottoy, Charisse Somers, Ellen De Roeck, Hanne Struyfs, Femke Soetewey, Jeroen Verhaeghe, et al. 2017. “The Cerebrospinal Fluid Aβ1-42/Aβ1-40 Ratio Improves Concordance with Amyloid-PET for Diagnosing Alzheimer’s Disease in a Clinical Setting.” Journal of Alzheimer’s Disease: JAD 60 (2): 561–76.

Palmqvist, Sebastian, Henrik Zetterberg, Niklas Mattsson, Per Johansson, Alzheimer’s Disease Neuroimaging Initiative, Lennart Minthon, Kaj Blennow, Mattias Olsson, Oskar Hansson, and Swedish BioFINDER Study Group. 2015. “Detailed Comparison of Amyloid PET and CSF Biomarkers for Identifying Early Alzheimer Disease.” Neurology 85 (14): 1240–49.

Pawlowski, Nick, S. Ira Ktena, Matthew C. H. Lee, Bernhard Kainz, Daniel Rueckert, Ben Glocker, and Martin Rajchl. 2017. “Dltk: State of the Art Reference Implementations for Deep Learning on Medical Images.” 10.48550/arXiv.1711.06853.

Petersen, R. C., P. S. Aisen, L. A. Beckett, M. C. Donohue, A. C. Gamst, D. J. Harvey, C. R. Jack Jr, et al. 2010. “Alzheimer’s Disease Neuroimaging Initiative (ADNI): Clinical Characterization.” Neurology 74 (3): 201–9.

Reith, Fabian H., Elizabeth C. Mormino, and Greg Zaharchuk. 2021. “Predicting Future Amyloid Biomarkers in Dementia Patients with Machine Learning to Improve Clinical Trial Patient Selection.” Alzheimer’s & Dementia: The Journal of the Alzheimer’s Association 7 (1). 10.1002/trc2.12212.

Reith, F., M. E. Koran, G. Davidzon, G. Zaharchuk, and Alzheimer′s Disease Neuroimaging Initiative. 2020. “Application of Deep Learning to Predict Standardized Uptake Value Ratio and Amyloid Status on F-Florbetapir PET Using ADNI Data.” AJNR. American Journal of Neuroradiology 41 (6): 980–86.

Richards, Danielle, and Marwan N. Sabbagh. 2014. “Florbetaben for PET Imaging of Beta-Amyloid Plaques in the Brain.” Neurology and Therapy 3 (2): 79–88.

Rieke, Johannes, Fabian Eitel, Martin Weygandt, John-Dylan Haynes, and Kerstin Ritter. 2018. “Visualizing Convolutional Networks for MRI-Based Diagnosis of Alzheimer’s Disease.” Understanding and Interpreting Machine Learning in Medical Image Computing Applications, 24–31.

Shaw, Leslie M., and John Q. Trojanowski. 2017. “Biomarker Core Report: Year1 ADNI3, Roche Elecsys Immunoassay Analyses of ADNI1/GO/2 CSF Samples.”

Shaw, Leslie M., Hugo Vanderstichele, Malgorzata Knapik-Czajka, Christopher M. Clark, Paul S. Aisen, Ronald C. Petersen, Kaj Blennow, et al. 2009. “Cerebrospinal Fluid Biomarker Signature in Alzheimer’s Disease Neuroimaging Initiative Subjects.” Annals of Neurology 65 (4): 403–13.

Spallazzi, Marco, Federica Barocco, Giovanni Michelini, Paolo Immovilli, Arens Taga, Nicola Morelli, Livia Ruffini, and Paolo Caffarra. 2019. “CSF Biomarkers and Amyloid PET: Concordance and Diagnostic Accuracy in a MCI Cohort.” Acta Neurologica Belgica 119 (3): 445–52.

Sperling, Reisa A., Jason Karlawish, and Keith A. Johnson. 2013. “Preclinical Alzheimer Disease-the Challenges Ahead.” Nature Reviews. Neurology 9 (1): 54–58.

Springenberg, Jost Tobias, Alexey Dosovitskiy, Thomas Brox, and Martin Riedmiller. 2014. “Striving for Simplicity: The All Convolutional Net,” December. 10.48550/arXiv.1412.6806.

Thal, Dietmar R., Udo Rüb, Mario Orantes, and Heiko Braak. 2002. “Phases of Aβ-Deposition in the Human Brain and Its Relevance for the Development of AD.” Neurology 58 (12): 1791–1800.

Weber, Christopher J., Maria C. Carrillo, William Jagust, Clifford R. Jack Jr, Leslie M. Shaw, John Q. Trojanowski, Andrew J. Saykin, et al. 2021. “The Worldwide Alzheimer’s Disease Neuroimaging Initiative: ADNI-3 Updates and Global Perspectives.” Alzheimer’s & Dementia: The Journal of the Alzheimer’s Association 7 (1). 10.1002/trc2.12226.

Wolk, David A., Zheng Zhang, Sanaa Boudhar, Christopher M. Clark, Michael J. Pontecorvo, and Steven E. Arnold. 2012. “Amyloid Imaging in Alzheimer’s Disease: Comparison of Florbetapir and Pittsburgh Compound-B Positron Emission Tomography.” Journal of Neurology, Neurosurgery, and Psychiatry 83 (9): 923–26.

