## Supplementary for "ArcheD, a residual neural network for prediction of cerebrospinal fluid amyloid-beta from amyloid PET images"

*Data used in preparation of this article were obtained from the Alzheimer's Disease Neuroimaging Initiative (ADNI) database (adni.loni.usc.edu). As such, the investigators within the ADNI contributed to the design and implementation of ADNI and/or provided data but did not participate in analysis or writing of this report. A complete listing of ADNI investigators can be found at: http://adni.loni.usc.edu/wp-content/uploads/how_to_apply/ADNI_Acknowledgement_List.pdf

Table S1. List of regions-of-interest (ROI) from Neuromorphometrics atlas

| **Name of the ROI** |
|---|
| 3rd Ventricle |
| 4th Ventricle |
| Brain Stem |
| Optic Chiasm |
| CSF |
| Accumbens Area |
| Amygdala |
| Caudate |
| Cerebellum Exterior (Cerebellum Gray Matter) |
| Cerebellum White Matter |
| Cerebral White Matter |
| Hippocampus |
| Inf Lat Vent |
| Lateral Ventricle |
| Pallidum |
| Putamen |
| Thalamus Proper |
| Ventral DC |
| Vessel |
| Basal Forebrain |
| Anterior cingulate gyrus (ACgG) |
| Anterior insula (AIns) |
| Anterior orbital gyrus (AOrG) |
| Angular gyrus (AnG) |
| Calcarine cortex (Calc) |
| Central operculum (CO) |
| Cuneus (Cun) |
| Entorhinal area (Ent) |
| Frontal operculum (FO) |
| Frontal pole (FRP) |
| Fusiform gyrus (FuG) |
| Gyrus rectus (GRe) |
| Inferior occipital gyrus (IOG) |
| Inferior temporal gyrus (ITG) |
| Lingual gyrus (LiG) |
| Lateral orbital gyrus (LOrG) |
| Middle cingulate gyrus (MCgG) |
| Medial frontal cortex (MFC) |
| Middle frontal gyrus (MFG) |
| Middle occipital gyrus (MOG) |
| Medial orbital gyrus (MOrG) |
| Postcentral gyrus medial segment (MPoG) |
| Precentral gyrus medial segment (MPrG) |
| Superior frontal gyrus medial segment (MSFG) |
| Middle temporal gyrus (MTG) |
| Occipital pole (OCP) |
| Occipital fusiform gyrus (OFuG) |
| Opercular part of the inferior frontal gyrus (OpIFG) |
| Orbital part of the inferior frontal gyrus (OrIFG) |
| Posterior cingulate gyrus (PCgG) |

Precuneus (PCu)
Parahippocampal gyrus (PHG)
Posterior insula (PIns)
Parietal operculum (PO)
Postcentral gyrus (PoG)
Posterior orbital gyrus (POrG)
Planum polare (PP)
Precentral gyrus (PrG)
Planum temporale (PT)
Subcallosal area (SCA)
Superior frontal gyru (SFG)
Supplementary motor cortex (SMC)
Supramarginal gyrus (SMG)
Superior occipital gyrus (SOG)
Superior parietal lobule (SPL)
Superior temporal gyrus (STG)
Temporal pole (TMP)
Triangular part of the inferior frontal gyrus (TrIFG)
Transverse temporal gyrus (TTG)

Table S2. Descriptive statistics of the study participants.

| | Clinical classification (*n*=1870) | | | | Biological classification (*n*=1865)* | | | |
|---|---|---|---|---|---|---|---|---|
| | SMC | CN | MCI | AD | A-T- | A-T+ | A+T- | A+T+ |
| Number of samples | 145 | 607 | 928 | 190 | 380 | 205 | 300 | 980 |
| Age (years, mean, SD) | 73 (5.4) | 75 (7.1) | 73 (7.5) | 74 (8.5) | 72 (7.4) | 75 (7.7) | 73 (7.4) | 74 (7.2) |
| Gender: Female (n, %) | 91 (62.8) | 345 (56.8) | 407 (43.9) | 73 (38.4) | 202 (53.2) | 109 (53.2) | 134 (44.7) | 468 (47.8) |
| Education (years, mean, SD) | 16.6 (2.4) | 16.8 (2.5) | 16.2 (2.6) | 15.7 (2.7) | 16.6 (2.5) | 16.4 (2.6) | 16.8 (2.5) | 16.1 (2.6) |
| Ethnicity (n,%): 1. African American | 4 (2.8) | 36 (5.9) | 21 (2.3) | 6 (3.2) | 18 (4.7) | 2 (1.0) | 16 (5.3) | 30 (3.1) |
| 2. White | 136 (93.8) | 550 (90.6) | 875 (94.3) | 178 (93.7) | 350 (92.1) | 197 (96.0) | 266 (88.7) | 922 (94.1) |
| 3. Others | 5 (3.4) | 21 (2.4) | 32 (3.4) | 6 (3.1) | 12 (3.2) | 6 (3.0) | 18 (6.0) | 27 (2.8) |
| Amyloid-beta (pg/mL, mean, SD) | 1124.8 (438.8) | 1249.5 (614.2) | 987.7 (544.0) | 604.3 (301.1) | 1527.9 (404.7) | 1841.8 (655.3) | 844.2 (329.2) | 752.1 (341.8) |
| Tau (pg/mL, mean, SD) | 234.1 (93.7) | 245.5 (92.3) | 287.3 (134.1) | 377.2 ( 150.1) | 194.6 (37.6) | 327.6 (69.9) | 171.7 (43.3) | 334.9 (136.7) |
| Phosphorylated Tau (pg/mL, mean, SD) | 26.8 (11.2) | 24.2 (10.1) | 28.3 (15.1) | 29.8 (33.1) | 16.6 (3.2) | 29.5 (7.3) | 16.0 (3.7) | 33.9 (19.2) |
| Apolipoprotein E (*APOE*) (n, %): 1. e2/e2 | – | – | – | 1 (0.5) | – | – | – | 1 (0.1) |
| 2. e2/e3 | 20 (13.8) | 57 (9.4) | 58 (6.2) | 6 (3.2) | 61 (16.1) | 16 (7.8) | 22 (7.3) | 42 (4.3) |
| 3. e2/e4 | 3 (2.1) | 8 (1.3) | 15 (1.6) | 2 (1.1) | 2 (0.5) | 4 (2.0) | 4 (1.3) | 18 (1.8) |
| 4. e3/e3 | 79 (54.5) | 365 (60.1) | 393 (42.3) | 50 (26.3) | 238 (62.6) | 132 (64.4) | 156 (52.0) | 356 (36.3) |
| 5. e3/e4 | 41 (28.3) | 135 (22.2) | 322 (34.7) | 76 (40.0) | 47 (12.4) | 43 (21.0) | 81 (27.0) | 403 (41.1) |
| 6. e4/e4 | 2 (1.4) | 19 (3.1) | 94 (10.1) | 37 (19.5) | 3 (0.8) | 1 (0.5) | 21 (7.0) | 127 (13.0) |
| 7. Unknown | – | 23 (3.8) | 46 (5.0) | 18 (9.5) | 29 (7.6) | 9 (4.4) | 16 (5.3) | 33 (3.4) |

Note: * Five samples were excluded due to inability to assign biological class in absence of phosphorylated tau measurements.

Amyloid-beta presents amyloid cerebrospinal fluid (CSF) measurement. In Apolipoprotein E (APOE) genotypes, e2 and e4 indicate the protective and risk alleles, respectively.

Abbreviation: SD – standard deviation; Clinical classes: AD – Alzheimer's disease; CN – cognitive normal; MCI – mild cognitive impairment; SMC – subjective memory concerns; Biological classes: 'A-T-' – negative amyloid and tau proteins CSF measurements based on the cut-off value; 'A-T+' – amyloid negative, tau positive; 'A+T-' – amyloid positive, tau negative; 'A+T+' – amyloid and tau positive.

Table S3. Comparison between linear and polynomial models' performance in fluid biomarker rescaling from INNO-BIA AlzBio3 system to Roche Elecsys.

| | Amyloid-beta | | |
|---|---|---|---|
| | $R^2$ train | $R^2$ test | Acc test |
| Linear regression | 0.759 | 0.689 | 0.915 |
| 3rd degree polynomial regression | 0.761 | 0.694 | 0.915 |

Abbreviation: $R^2$ – coefficient of determination; Acc – accuracy.

Table S4. Mean relevance values for brain regions and 95% bootstrapped confidence intervals (CI). Values were multiplied by 1000.

| Brain region | Mean | Lower 95% CI | Upper 95% CI |
|---|---|---|---|
| **Cerebellum white matter** | 0.252 | 0.248 | 0.256 |
| **Cerebral white matter** | 0.207 | 0.205 | 0.208 |
| **Brain stem** | 0.134 | 0.133 | 0.136 |
| **Subcortical areas** | 0.112 | 0.110 | 0.113 |
| **Temporal lobe** | 0.106 | 0.105 | 0.108 |
| **Parietal lobe** | 0.105 | 0.104 | 0.107 |
| **Occipital lobe** | 0.099 | 0.098 | 0.100 |
| **Vessels** | 0.099 | 0.095 | 0.103 |
| **Frontal lobe** | 0.091 | 0.089 | 0.092 |
| **Cerebellum Gray Matter** | 0.089 | 0.088 | 0.090 |
| **Limbic lobe** | 0.046 | 0.045 | 0.047 |
| **Basal forebrain** | 0.036 | 0.034 | 0.038 |
| **Ventricles** | 0.014 | 0.013 | 0.015 |
| **Optic chiasm** | 0.012 | 0.012 | 0.014 |

Table S5. Comparison of brain regions' relevance values distribution within clinical and biological classifications.

| | AD vs CN (*q; d*) | MCI vs CN | SMC vs CN | A+T+ vs A-T- | A+T- vs A-T- | A-T+ vs A-T- |
|---|---|---|---|---|---|---|
| **Cerebellum Gray Matter** | >0.01; 0.294 | >0.01; 0.047 | >0.01; -0.012 | 1.009e-05; 0.321 | >0.01; 0.162 | >0.01; -0.073 |
| **Cerebellum White Matter** | >0.01; -0.222 | >0.01; -0.034 | >0.01; -0.049 | 1.798e-03; 0.214 | >0.01; 0.104 | >0.01; 0.045 |
| **Cerebral White Matter** | 3.997e-20; -0.976 | 2.943e-08; -0.358 | >0.01;0.114 | 1.723e-48; -1.138 | 2.152e-12; -0.720 | >0.01; 0.025 |
| **Vessel** | >0.01; 0.141 | >0.01; 0.062 | >0.01; -0.067 | >0.01; -0.007 | >0.01; 0.049 | >0.01; 0.036 |
| **Basal Forebrain** | 6.568e-09; 0.840 | 5.363e-09; 0.344 | >0.01; 0.242 | 1.545e-21; 0.542 | >0.01; 0.297 | >0.01; -0.015 |
| **Brain Stem** | >0.01; 0.298 | 4.1346e-05; 0.266 | >0.01; -0.121 | 2.108e-09; 0.406 | 2.194e-03; 0.337 | >0.01; 0.027 |
| **Frontal lobe** | 3.378e-10; 0.785 | 4.184e-06; 0.288 | >0.01; 0.053 | 7.759e-31; 0.761 | 1.539e-04; 0.446 | >0.01; 0.128 |
| **Limbic lobe** | 5.271e-06; 0.610 | 2.335227e-05; 0.243 | >0.01; -0.008 | 9.371e-08; 0.321 | >0.01; 0.230 | >0.01; -0.047 |
| **Occipital lobe** | 3.853e-15; 0.970 | 3.936e-10; 0.400 | >0.01; -0.115 | 1.723e-48; 1.017 | 4.793e-06; 0.514 | >0.01; 0.061 |
| **Optic Chiasm** | >0.01; 0.044 | >0.01; 0.051 | >0.01; 0.008 | >0.01; -0.079 | >0.01; -0.020 | >0.01; -0.194 |
| **Parietal lobe** | 9.558e-09; 0.712 | 3.551e-05; 0.259 | >0.01; 0.006 | 6.500e-31; 0.764 | 1.767e-03; 0.371 | >0.01; 0.121 |
| **Subcortical** | 4.434e-16; 0.909 | 9.215e-05; 0.260 | >0.01; 0.011 | 1.952e-20; 0.650 | 5.353e-04; 0.383 | >0.01; -0.008 |
| **Temporal lobe** | 3.333e-21; 1.169 | 4.193e-10; 0.393 | >0.01; -0.026 | 6.446e-102; 1.493 | 1.991e-12; 0.831 | >0.01; 0.184 |
| **Ventricles** | >0.01; 0.227 | >0.01; 0.142 | >0.01; 0.020 | >0.01; 0.170 | >0.01; -0.050 | >0.01; 0.018 |

Abbreviation: *q* – FDR-corrected t-test *p*-value; *d* – Cohen's *d* coefficient.

Table S6. Mean relevance values for cortical regions and cerebellum exterior and bootstrapped 95% confidence intervals (CI) separately for all clinical and biological classes. Values were multiplied by 1000.

| | **Clinical classes (mean, [Lower 95% CI, Upper 95% CI])** | | | | **Biological classes (mean, [Lower 95% CI, Upper 95% CI])** | | | |
|---|---|---|---|---|---|---|---|---|
| **Brain region** | **CN** | **SMC** | **MCI** | **AD** | **A-T-** | **A-T+** | **A+T-** | **A+T+** |
| **Temporal lobe** | 0.098, [0.095, 0.100] | 0.097, [0.093, 0.101] | 0.108, [0.106, 0.110] | 0.126, [0.122, 0.131] | 0.087, [0.085, 0.089] | 0.091, [0.089, 0.093] | 0.104, [0.100, 0.108] | 0.124, [0.122, 0.126] |
| **Parietal lobe** | 0.101, [0.099, 0.103] | 0.101, [0.098, 0.104] | 0.106, [0.105, 0.108] | 0.116, [0.112, 0.120] | 0.097, [0.095, 0.098] | 0.099, [0.097, 0.101] | 0.104, [0.100, 0.108] | 0.113, [0.112, 0.115] |
| **Occipital lobe** | 0.093, [0.091, 0.095] | 0.091, [0.088, 0.094] | 0.101, [0.100, 0.103] | 0.112, [0.108, 0.116] | 0.089, [0.087, 0.090] | 0.090, [0.088, 0.091] | 0.098, [0.095, 0.101] | 0.109, [0.108, 0.111] |
| **Frontal lobe** | 0.086, [0.084, 0.088] | 0.087, [0.084, 0.090] | 0.092, [0.090, 0.093] | 0.101, [0.097, 0.105] | 0.083, [0.081, 0.084] | 0.085, [0.083, 0.086] | 0.091, [0.088, 0.094] | 0.098, [0.096, 0.100] |
| **Cerebellum Gray Matter** | 0.088, [0.086, 0.090] | 0.088, [0.084, 0.091] | 0.089, [0.087, 0.090] | 0.095, [0.090, 0.099] | 0.086, [0.083, 0.088] | 0.084, [0.082, 0.087] | 0.089, [0.086, 0.094] | 0.093, [0.091, 0.095] |

Abbreviation: Clinical classes: AD – Alzheimer's disease; CN – cognitive normal; MCI – mild cognitive impairment; SMC – subjective memory concerns; Biological classes: 'A-T-' – negative amyloid and tau proteins CSF measurements based on the cut-off value; 'A-T+' – amyloid negative, tau positive; 'A+T-' – amyloid positive, tau negative; 'A+T+' – amyloid and tau positive.

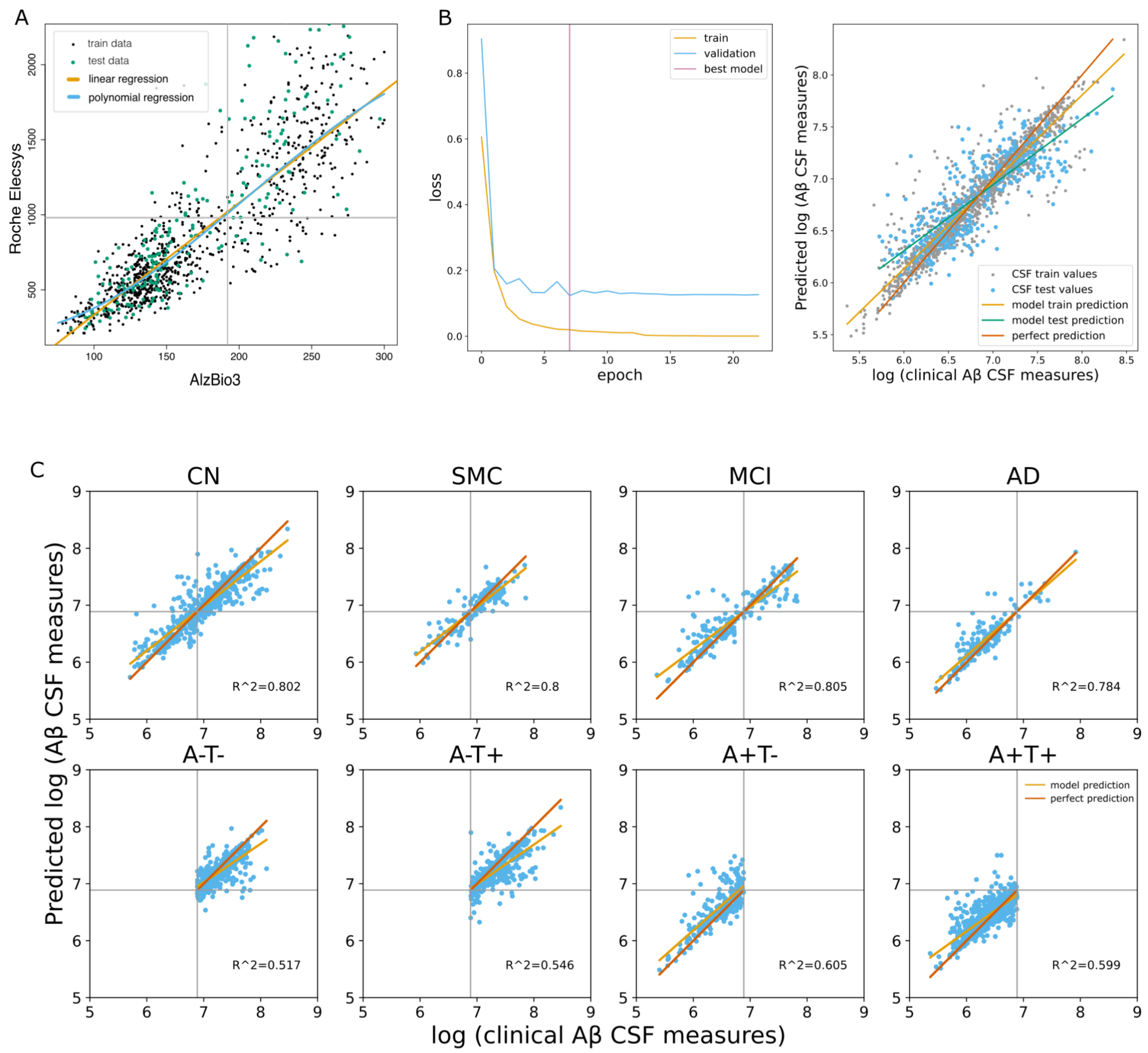


Figure S1. (A) Linear (orange line) and 3rd degree polynomial (blue line) regression for fluid amyloid rescaling from INNO-BIA AlzBio3 system to Roche Elecsys. Gray lines present cut-off values for Aβ CSF measurement. (B) (left) Logarithmic model loss (MSE) for training and validation datasets. (right) Comparison of clinically measured Aβ CSF and predicted by the model. Red line - case when the model predicts labels ideally. Green line - linear relation between real and predicted values of the test dataset. Orange line - linear relation between real and predicted values of the training dataset. (C) Comparison of clinically measured Aβ CSF and predicted by the model for clinical and biological classes. Red line - case when the model predicts labels ideally. Orange line - linear relation between real and predicted amyloid CSF values for a concrete subgroup. Abbreviation: Aβ – amyloid-beta; CSF – Cerebrospinal fluid; Clinical classes: AD – Alzheimer's disease; CN – cognitive normal; MCI – mild cognitive impairment; SMC – subjective memory concerns; Biological classes: 'A-T-' – negative amyloid and tau proteins CSF measurements based on the cut-off value; 'A-T+' – amyloid negative, tau positive; 'A+T-' – amyloid positive, tau negative; 'A+T+' – amyloid and tau positive.

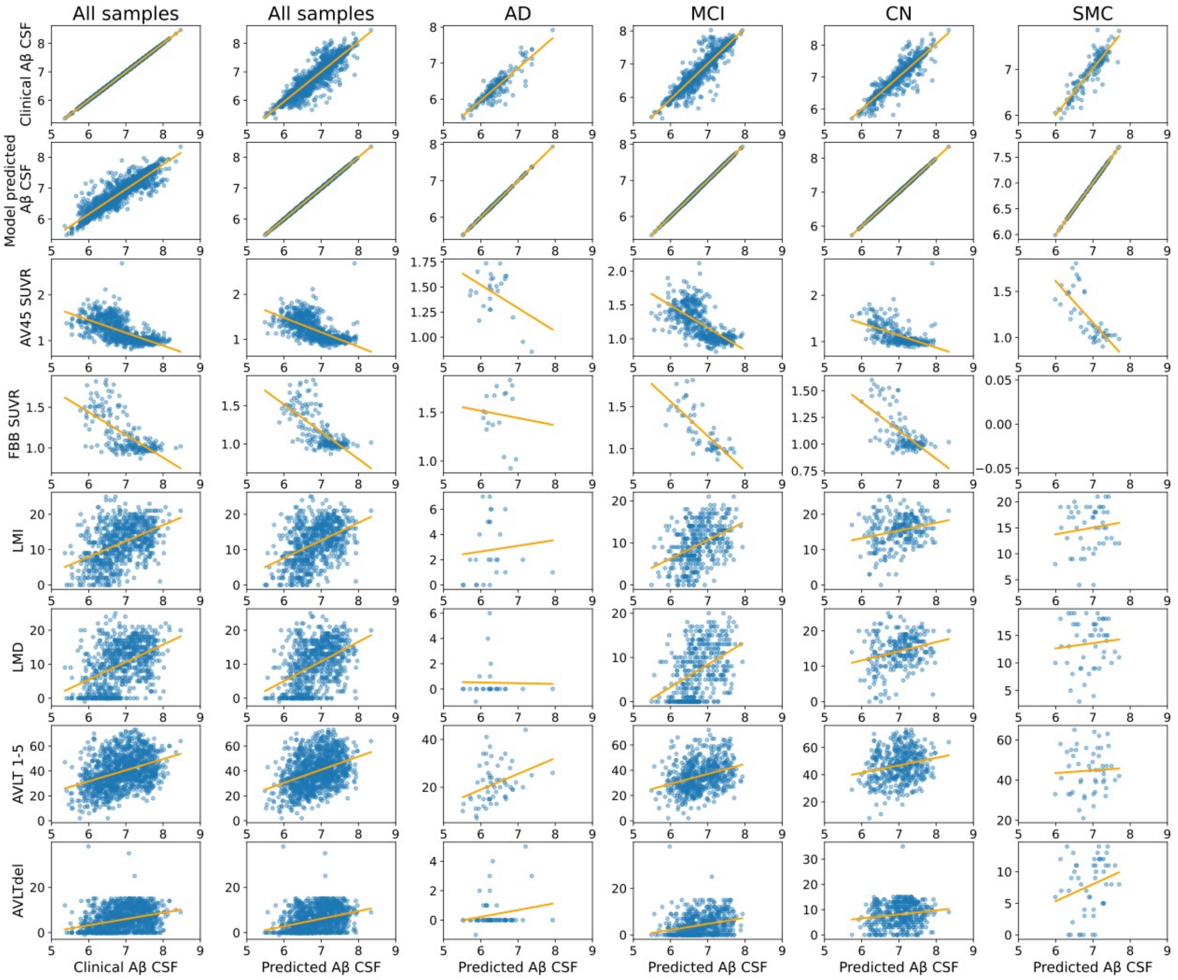


Figure S2. Correlation scatter plots for AD hallmark biomarker. Aβ - amyloid-beta; CSF - Cerebrospinal fluid; AD - Alzheimer's disease; CN - cognitively normal; MCI - mild cognitive impairment; SMC - subjective memory concerns; SUVR - standardized uptake value ratio; LMI - immediate recall measurements of Logical Memory (LM) II subtests of the Wechsler Memory Scale-Revised; LMD - delayed recall measurements of LM; AVLT 1-5 - immediate (total words in trials 1-5) recall measurements of the Rey Auditory Verbal Learning Test (AVLT); AVLTdel - delayed recall measurements of the AVLT. Orange line presents a linear relation between parameters.

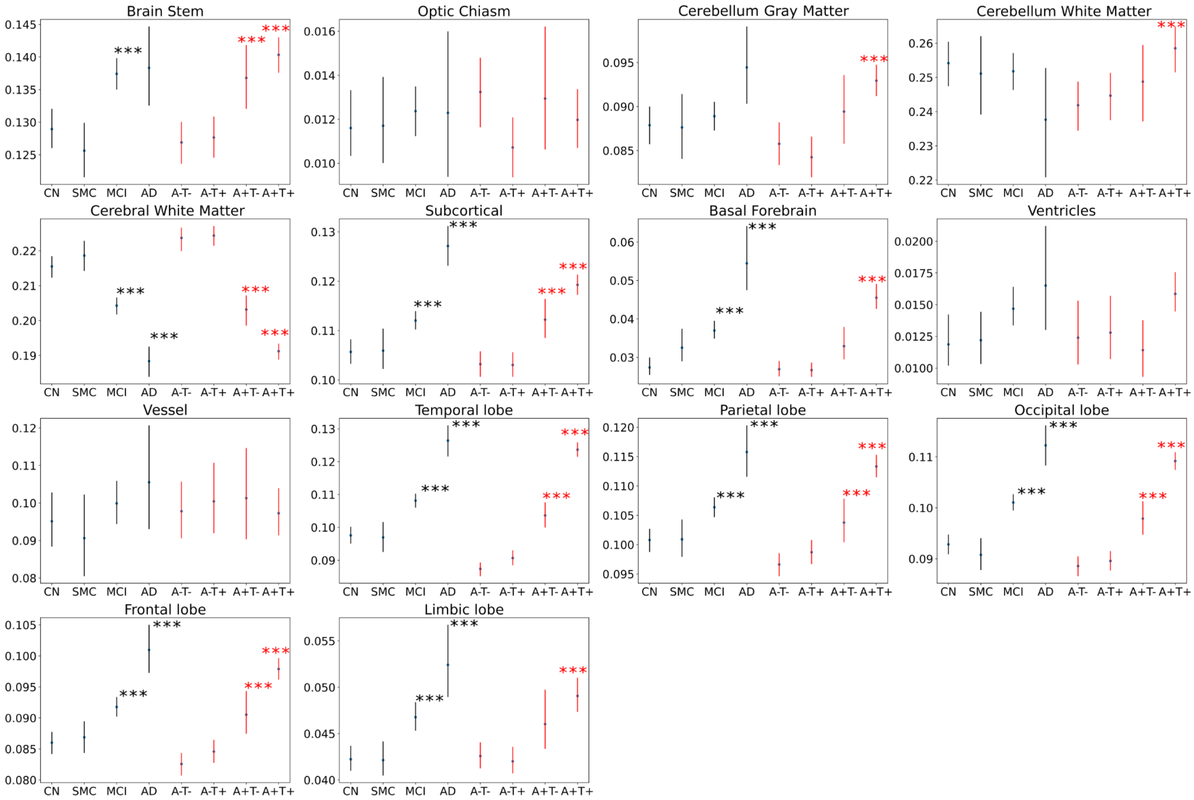


Figure S3. Comparison of mean relevance values per brain region within clinical and biological classifications. Cognitive normal (CN) group was used as control for clinical classification. A-T- was used as control for biological classification.

Abbreviation: Clinical classes: AD – Alzheimer's disease; CN – cognitive normal; MCI – mild cognitive impairment; SMC – subjective memory concerns; Biological classes: 'A-T-' – negative amyloid and tau proteins CSF measurements based on the cut-off value; 'A-T+' – amyloid negative, tau positive; 'A+T-' – amyloid positive, tau negative; 'A+T+' – amyloid and tau positive.

*** – $q$<0.01.
